# Biofilm formation in *Streptococcus suis*: *In vitro* impact of serovar and assessment of coinfections with other porcine respiratory disease complex bacterial pathogens

**DOI:** 10.1101/2024.06.26.600819

**Authors:** Rubén Miguélez-Pérez, Oscar Mencía-Ares, César B. Gutiérrez-Martín, Alba González-Fernández, Máximo Petrocchi-Rilo, Mario Delgado-García, Sonia Martínez-Martínez

## Abstract

*Streptococcus suis* is a worldwide pathogen that impacts swine industry, causing severe clinical signs in postweaning piglets, including meningitis and arthritis. Biofilm formation is a major virulence mechanism in *S. suis*, enhancing its persistence and resistance. Here, we assessed the *in vitro* biofilm formation of 240 *S. suis* isolates from Spanish swine farms and evaluated the effects of serovars (SVs) and coinfections with other porcine respiratory disease complex (PRDC) pathogens. Our study revealed significant heterogeneity in biofilm formation among *S. suis* SVs. Notably, SV2 exhibited the lowest biofilm formation, contrasting with the high biofilm-forming capacities of SV1, SV7, and SV9. Virulence factors *epf*, *mrp*, and *sly* were associated (*p* < 0.05) with reduced biofilm formation. Other PRDC pathogens, including *Actinobacillus pleuropneumoniae*, *Glaesserella parasuis*, and *Pasteurella multocida*, formed biofilms, though generally less robust than those of *S. suis* (except for SV2), contrasting the high biofilm formation of *Staphylococcus hyicus*. Coinfections demonstrated enhanced biofilm formation in mixed cultures of *S. suis*, particularly with *P. multocida*. Other coinfections revealed variable results in pathogen interactions, suggesting the potential of biofilms for increased persistence and pathogenicity in coinfections. In conclusion, this study underscores the importance of serovar-specific differences in biofilm formation among *S. suis* isolates, with significant implications for pathogenicity and persistence. The heterogeneous biofilm formation observed in coinfections with other PRDC pathogens reveals a complex interplay that could exacerbate disease severity. These findings provide a foundation for further research on biofilm mechanisms to mitigate the impact of PRDC in the swine industry.

## Introduction

*Streptococcus suis* constitutes a worldwide hazard, not only because of its impact in the swine industry, but also as a zoonotic pathogen (1). Within pig production, *S. suis* mostly affects postweaning piglets between four and ten weeks of age. Once the upper respiratory tract is colonized, the host usually develops an asymptomatic carriage, eventually leading to the development of invasive disease, causing severe clinical signs, including arthritis, meningitis, endocarditis, septicaemia, and ultimately, sudden death (2).

*S. suis* exhibits significant genetic and phenotypic heterogeneity, even among strains of the same serovar (SV). Currently, there are 29 well-defined SVs (1–19, 21, 23–25, 27– 31, and 1/2) based on the antigenicity of the capsular polysaccharides (3). Over a hundred virulence factors (VFs) have been described for *S. suis* in the literature (4). However, despite the critical role of some of them, no specific VF has been deemed necessary for the disease (5). Among them, we could highlight the particular role of adhesins and cell surface factors, such as the muramidase-released protein (MRP), the extracellular factor (EF), and the glyceraldehyde-3-phosphate dehydrogenase (GAPDH); toxins, as the suilysin; and the S-ribosylhomocysteinase (LuxS), an interspecies quorum sensing related enzyme (6).

Biofilm formation is a major pathogenic factor in *S. suis*, facilitating its establishment in pig tissues (7). Biofilms are inter- or intraspecific communities of bacteria enclosed in a self-produced extracellular matrix, adhering to biotic or abiotic surfaces (8). They have been linked to increased resistance to antimicrobial agents, environmental stress, and host immune system, contributing to the chronicity of infections (9). Despite research on *S. suis* biofilms has been heavily promoted since its first description in 2007 (10), the understanding of its formation mechanisms remain superficial (7). Moreover, little is known about the role of *S. suis* on biofilm formation in coinfections with other bacterial pathogens directly or indirectly involved in the porcine respiratory disease complex (PRDC), a multifactorial syndrome affecting the respiratory system of postweaning piglets (11).

For all these reasons, the aim of this study was the assessment of the *in vitro* biofilm formation and the characterization of VFs of a selection of 240 *S. suis* recovered from Spanish pig farms belonging to different SVs, together with the evaluation of the *in vitro* biofilm formation capacity of a selection of bacterial pathogens frequently involved in the PRDC, alone and in coinfection with *S. suis*.

## Materials and methods

### Bacterial isolates and growth conditions

A wide range of different bacterial isolates have been used in this study. First, 240 *S. suis* isolates belonging to 16 different SVs (*i.e*., 1, 2, 3, 4, 5, 7, 8, 9, 10, 12, 16, 17, 18, 19, 21 and 31) and isolated from three anatomic regions: central nervous system (CNS), lungs and joints. As for the other bacteria, a total of 35 *Glaesserella parasuis*, 31 *Staphylococcus hyicus*, 20 *Pasteurella multocida* and 12 *Actinobacillus pleuropneumoniae* isolates were tested. These isolates were recovered from clinical cases from Spanish swine farms collected between 2020 and 2024, and further included in the strain collection of the BACRESPI research group at the Animal Health Department of the University of León (Spain).

*S. suis* isolates were cultured on Todd-Hewitt broth (THB) agar (Condalab, Spain) supplemented with 5% (v/v) fetal bovine serum (FBS) (Gibco, USA) and grown at 37LJ for 24 h under aerophilic conditions. The rest of the bacterial isolates were cultured on chocolate agar plates with Vitox (Oxoid, UK). *S. hyicus* and *P. multocida* were incubated at 37°C for 24 h under aerophilic conditions, while *G. parasuis* and *A. pleuropneumoniae* were incubated at 37°C for 48 h under microaerophilic conditions.

### Molecular characterization of bacterial isolates

Serotyping and virulence-related characterization of the different bacterial species was accomplished via polymerase chain reaction (PCR). *S. suis* characterization was based on the protocol described by Petrocchi *et al*. (12), which included various sets of a multiplex PCRs to identify all SVs, and five monoplex PCRs for virulence-associated genes, including *epf*, *gapdh*, *luxS*, *mrp*, and *sly*.

For *S. hyicus*, a multiplex PCR was used for detection of genes encoding exfoliative toxins ExhA, ExhB, ExhC and ExhD, as described by Andresen & Ahrens (13). Six of the most relevant virulence-associated genes of *P. multocida* were detected via PCR, as specified by Ewers *et al.* (14), including *hgbA*, *ompH*, *nanH*, *sodA*, *oma87*, and *pfhA*. Another multiplex PCR was used to differentiate the serogroups of *cap* gene A, B, D, E and F, as described by Townsend *et al.* (15). In *G. parasuis* isolates, a PCR was utilized to classify isolates into virulent or non-virulent based on *vtaA* genes, as described by Galofré-Milá et al. (16). Regarding to *A. pleuropneumoniae*, a monoplex PCR was utilized to classify isolates based on the iron-repressible outer-membrane protein 1, following the protocol from de la Puente-Redondo et. al. (17).

### Biofilm formation assay by single and mixed cultures

Biofilm formation of all the different isolates was quantified by crystal violet staining, following an archetypical biofilm formation protocol previously described (18) with slight modifications. Briefly, for *S. suis*, *S. hyicus* and *P. multocida*, a single colony was inoculated into 96-well polystyrene microfiber cell culture-treated plates (Corning Incorporated, USA) containing 200 µL of THB supplemented with 5% FBS. In the case of mixed cultures, a single colony of *S. suis* was inoculated, followed by the inoculation of the respective single colony of either *S. hyicus* or *P. multocida*. For both single colony and mixed cultures involving these pathogens, plates were incubated for 24 h under aerophilic conditions. For plates involving either single *A. pleuropneumoniae* and *G. parasuis* isolates or mixed cultures with *S. suis*, THB was supplemented with 5% FBS, 0.5% glucose (v/v) and 20 mg/mL NAD, with an incubation of 48 h under microaerophilic conditions.

In either case, following incubation, the culture medium and unattached bacteria were aspirated to remove them. The formed biofilms were stained with 100 μL of 2% crystal violet for 30 minutes, washed three times with distilled water, and dried at 37°C for 15 minutes. To release the dye, 100 μL of 95% ethanol was added and the plates were briefly agitated. The absorbance of the biofilm biomass was quantified at 595 nm (A595). All assays were conducted in triplicate to ensure reliability of the results. The final optical density (OD) value of each isolate was expressed as the mean of the three measurements subtracting the average OD of the negative control (difference with the control, DC), to lessen the possible unevenness in absorbance quantification.

### Scanning electron microscopy (SEM) of biofilms

The SEM assay was conducted on a selection of four *S. suis* isolates belonging to the four main SVs (SV1, SV2, SV7, and SV9), and a selection of two mixed infections of each bacterial pathogen evaluated with *S. suis*. It was performed following a previously published method (19) with minor adjustments. Briefly, Thermanox Plastic Coverslips (13 mm in diameter and 0.13 mm in nominal thickness; NUNC, USA) were used as adhesion carriers for the biofilm, which were positioned into the bottom of the wells in 24-well polystyrene microfiber cell culture-treated plates (Corning Incorporated, USA).

Independently from the culture, and after its pertinent incubation time (with all medium, supplement volumes and number of colonies proportionately extrapolated), samples were fixed in 2.5% glutaraldehyde in phosphate buffer (PBS, 0.1 M, pH 7.4) at 4 °C for 12 h, rinsed three consecutive times with PBS, post-fixed with 1% osmium tetroxide in PBS for 45 min in the dark and rinsed again three times with PBS. Samples were dehydrated in graded ethanol series (30%, 50%, 70%, 90%, 3 x 96%, and 3 x 100%, each for 10 min), dried by the critical point method (CPD300, Leica, Austria), mounted on aluminium stubs with conducting carbon ribbon and sputter-coated with gold-palladium (ACE200, Leica, Austria). The samples were observed under a Jeol JSM-6840LV scanning electron microscope (Jeol, Japan) at 5 kV. Samples were first approached via a broad sweep visualization, to then further scrutinize representative areas to obtain images at either 2500×, 5000× or 10000× magnification.

### Data analysis and results visualization

Databases were created in several Excel sheets (Microsoft Office). The first database, used for *S. suis* characterization and biofilm formation evaluation, included *S. suis* ID, anatomic location (lung, joint, CNS), SV, presence/absence of VF (*epf*, *mrp*, *sly*, *luxS*, and *gapdh*), and biofilm formation. Biofilm formation was expressed numerically as DC and categorized based on DC value into low (DC ≤ 2), medium (2 > DC ≤ 3), and high (DC > 3), as previously described (20).

A second database included the four most clinically relevant *S. suis* SVs (SV1, SV2, SV7, and SV9) and four additional bacterial pathogens also frequently associated with *S. suis* infections (*S. hyicus*, *P. multocida*. *G. parasuis* and *A. pleuropneumoniae*). In addition to the information from the first database, specific details were recorded for each microorganism: the VFs (*exhA*, *exhB*, *exhC*, and *exhD*) for *S. hyicus*; the VFs (*hgbA*, *ompH*, *sodA*, *pfhA*, and *oma87*) and capsular type (A, B, D, E, and F) for *P. multocida*; the virulence (virulent/non-virulent isolate) for *G. parasuis*; and the SV for *A. pleuropneumoniae*. Coinfections were studied in a specific database for each pathogen pair, including all previous information along with biofilm formation in each coinfection.

Statistical analyses of DC values were conducted using non-parametric methods due to the non-normal distribution of the data. Differences were assessed using the Wilcoxon rank-sum test, with *p*-values adjusted following the Benjamini & Hochberg method, and significance established at *p* < 0.05. Analyses for each bacterial pathogen were performed initially for all isolates, and further itemized by SV, anatomic location, or any other specific variables. *S. suis* isolates belonging to SV18 (*n* = 1), SV19 (*n* = 1), SV31 (*n* = 1), and non-typified isolates (n = 7) were excluded from the statistical analysis when itemizing by SV due to their low frequency.

Ordination of *S. suis* isolates based on their VF composition was estimated using the Jaccard distance matrix and analyzed by principal component analysis (PCA). The two main dimensions for the principal components were characterized. The effect of biofilm formation capacity (low, medium, high) was determined by permutational multivariate analysis of variance (PERMANOVA) using distance matrices with *adonis2* function (pairwise adonis).

For the evaluation of coinfections, a selection of four isolates belonging to the four most clinically relevant *S. suis* SVs – SV1 (ID 990), SV2 (ID 1001), SV7 (ID 998), and SV9 (ID 969) – was made. These isolates were compared with a selection of ten random isolates from *S. hyicus*, *P. multocida*, *G. parasuis* and *A. pleuropneumoniae.* All analyses were performed within each *S. suis* SV and bacterial pathogen. To determine the effect of the *S. suis* SV on the biofilm formation of each bacterial pathogen, the analysis compared the DC average of the bacterial pathogen with the DC average of the coinfection with the specific *S. suis* SV. Similarly, to determine the effect of the bacterial pathogen on the biofilm production of the *S. suis* SV, the analysis compared the DC average of the specific *S. suis* with the DC of the coinfection with the pathogen.

All analyses were conducted using R version 4.3.2 (2023-10-31 ucrt) (21). Plots were produced using the *ggplot2* package (22) and further modified using the software *Inkscape* version 1.3.2 (https://inkscape.org/).

## Results

### Characterization of *Streptococcus suis* clinical isolates based on their virulence, serotyping, and anatomic location

Most isolates carried at least one VF, with frequencies of all VFs exceeding 60%. The most common VF was *luxS* (90%), followed by *gapdh* (80%), *epf* (64.6%), *mrp* (64.2%) and *sly* (62.5%). When evaluating the combinations of all VFs, 27 distinct patterns were observed. The most frequent combination was the presence of all genes (33.8%), followed by the combination of *epf*, *sly*, *lux*S and *gapdh*, and the combination of *luxS* and *gapdh*, each representing 8.3% of all *S. suis* isolates. A detailed description of VF patterns, both individually and by SV, is available in additional files (see Additional files 1 and 2).

Analyzing the isolates by SV, we found significant associations between certain SVs and VFs. Notably, there was a positive association of *epf*, *sly* and *mrp* with SVs 1, 2 and 9 (*p <* 0.001), and a negative association of *epf* and *sly* with SV7 (*p* < 0.05). Both *luxS* and *gapdh* were significantly more frequent in most isolates, regardless of SV, due to their high prevalence in the isolates of the study. A detailed description of these combinations is available in an additional file (see Additional file 3). Regarding *S. suis* anatomic location, we could determine that the only significant finding was a negative association between SV2 and lungs (*p <* 0.01), as most SV2 isolates were recovered from CNS.

### Influence of the serovar and the virulence factors in the biofilm formation capacity of *Streptococcus suis*

The SV deeply influenced the capacity of *S. suis* to produce biofilm (Figure 1A, Table 1). The most remarkable finding was the significantly lower biofilm formation of SV2 (DC = 1.77 ± 0.46) compared to most of the SVs commonly isolated in swine streptococcal infections (*p <* 0.05), demonstrating its low biofilm formation capacity (DC ≤ 2), as it was observed in SEM images (Figure 1B). In contrast, other clinically relevant *S. suis* SVs, such as SV7 (DC = 6.98 ± 2.33) and SV9 (DC = 5.77 ± 1.56), showed a strong biofilm formation capacity (Figure 1B). Overall, the mean biofilm formation for all SVs, except SV2, was categorized as strong (DC > 3). A detailed summary of the comparisons of biofilm formation capacity among SVs is available in an additional file (see Additional file 4). No significant association was observed between the anatomic locations of *S. suis* and its biofilm formation capacity.

**Figure 1.**
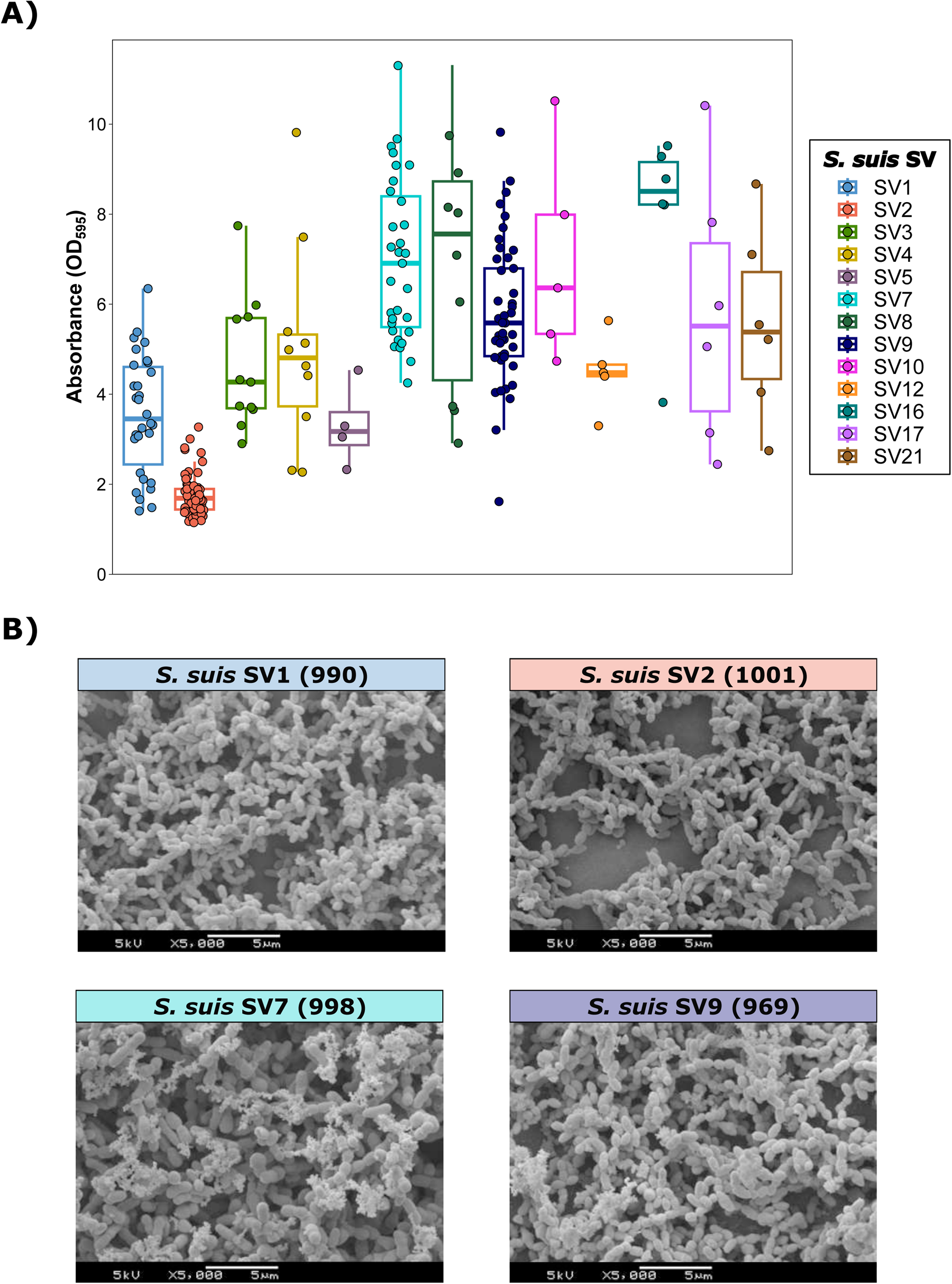
Biofilm formation of *Streptococcus suis* isolates recovered from Spanish swine farms. A) Boxplots illustrating biofilm formation categorized by *S. suis* serovar (SV). Biofilm formation for each isolate was quantified as the difference from the negative control (DC) in absorbance (OD_595_). Each *S. suis* isolate is represented by a dot with horizontal jitter for visibility. The horizontal box lines represent the first quartile, the median, and the third quartile. Whiskers extend to 1.5 interquartile range. B) Scanning electron microscope (SEM) images of biofilm formation in *S. suis* isolates belonging to the four main SVs (ID): *S. suis* SV1 (990); *S. suis* SV2 (1001); *S. suis* SV7 (998); and *S. suis* SV9 (969).

**Table 1.** Biofilm formation in *Streptococcus suis* isolates recovered from Spanish swine farms.

| *S. suis* SV | Number of isolates (*n*) | Biofilm formation (mean ± SD) | Biofilm formation (Degree) |
| --- | --- | --- | --- |
| SV1 | 30 | 3.54 ± 1.32 | High |
| SV2 | 61 | 1.77 ± 0.46 | Low |
| SV3 | 11 | 4.64 ± 1.46 | High |
| SV4 | 10 | 4.99 ± 2.28 | High |
| SV5 | 4 | 3.30 ± 0.92 | High |
| SV7 | 31 | 6.98 ± 1.78 | High |
| SV8 | 10 | 6.96 ± 2.82 | High |
| SV9 | 45 | 5.77 ± 1.56 | High |
| SV10 | 5 | $6.99 \pm 2.33$ | High |
| SV12 | 5 | $4.49 \pm 0.83$ | High |
| SV16 | 6 | $7.97 \pm 2.10$ | High |
| SV17 | 6 | $5.81 \pm 2.97$ | High |
| SV21 | 6 | $5.55 \pm 2.12$ | High |

The analysis of the association between biofilm formation and VF composition (Figure 2A) revealed that isolates carrying *epf* (*p <* 0.0001), *mrp* (*p <* 0.01), and *sly* (*p* < 0.01) had a lower biofilm formation capacity than those not carrying these genes. No significant differences were observed for *gapdh* and *luxS*. When evaluating the association between the degree of biofilm formation and VF combinations, we observed a slight (R^2^ = 0.03) but significant association (*p <* 0.001) using PERMANOVA analysis (Figure 2B), with a slightly lower variation in VF composition among isolates with low biofilm formation capacity (DC ≤ 2).

**Figure 2.**
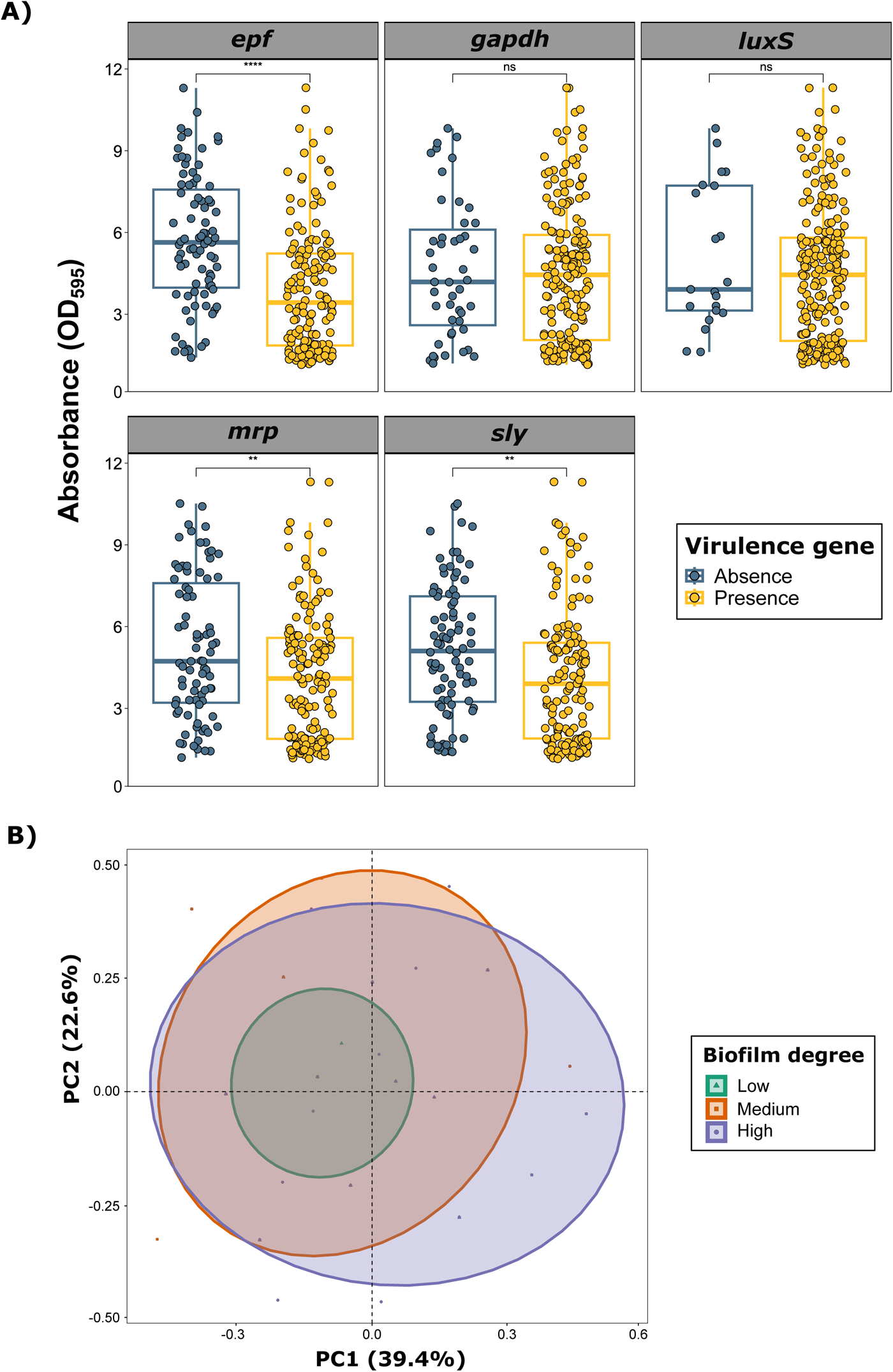
Impact of virulence factor (VF) genes on biofilm formation in *Streptococcus suis* isolates recovered from Spanish swine farms. A) Boxplots illustrating the quantitative biofilm formation of *S. suis*, comparing the presence or absence of VF genes. Quantification was performed as the difference from the negative control (DC) in absorbance (OD_595_). Each *S. suis* isolate is represented by a dot with horizontal jitter for visibility. The horizontal box lines represent the first quartile, the median, and the third quartile. Whiskers extend to 1.5 times the interquartile range. Differences between groups were evaluated using the Wilcoxon rank-sum test. B) Principal component analysis (PCA) of the five evaluated VF genes, showing grouping based on biofilm formation degrees.

When itemizing the information by SV, the results varied. For instance, SV1 isolates carrying the *epf* gene exhibited significantly higher biofilm formation (*p* < 0.01), while SV2 isolates with the *epf* gene (*p* < 0.05), SV9 isolates with the *sly* gene (*p <* 0.01), and SV1 isolates with the *gapdh* gene (*p* < 0.05) showed lower biofilm formation capacities.

### Biofilm formation in swine bacterial pathogens associated with *Streptococcus suis* infections

A huge variability in biofilm production was observed among those pathogens frequently associated with *S. suis* infections (Figure 3). *S. hyicus* exhibited the highest biofilm formation capacity (DC = 13.91 ± 8.51) among all tested species and *S. suis* SVs (*p <* 0.001), despite notable variability among isolates. It was followed by *S. suis* SV7 (DC = 7.02 ± 1.80), SV9 (DC = 5.77 ± 1.56), and SV1(DC = 3.54 ± 1.32). *P. multocida* (DC = 2.83 ± 2.01), and *G. parasuis* (DC = 2.66 ± 0.88) were categorized as medium biofilm producers (DC > 2 ≤ 3). *A. pleuropneumoniae* (DC = 1.76 ± 0.51) and *S. suis* SV2 (DC = 1.76 ± 0.45) exhibited the lowest biofilm formation capacities.

**Figure 3.**
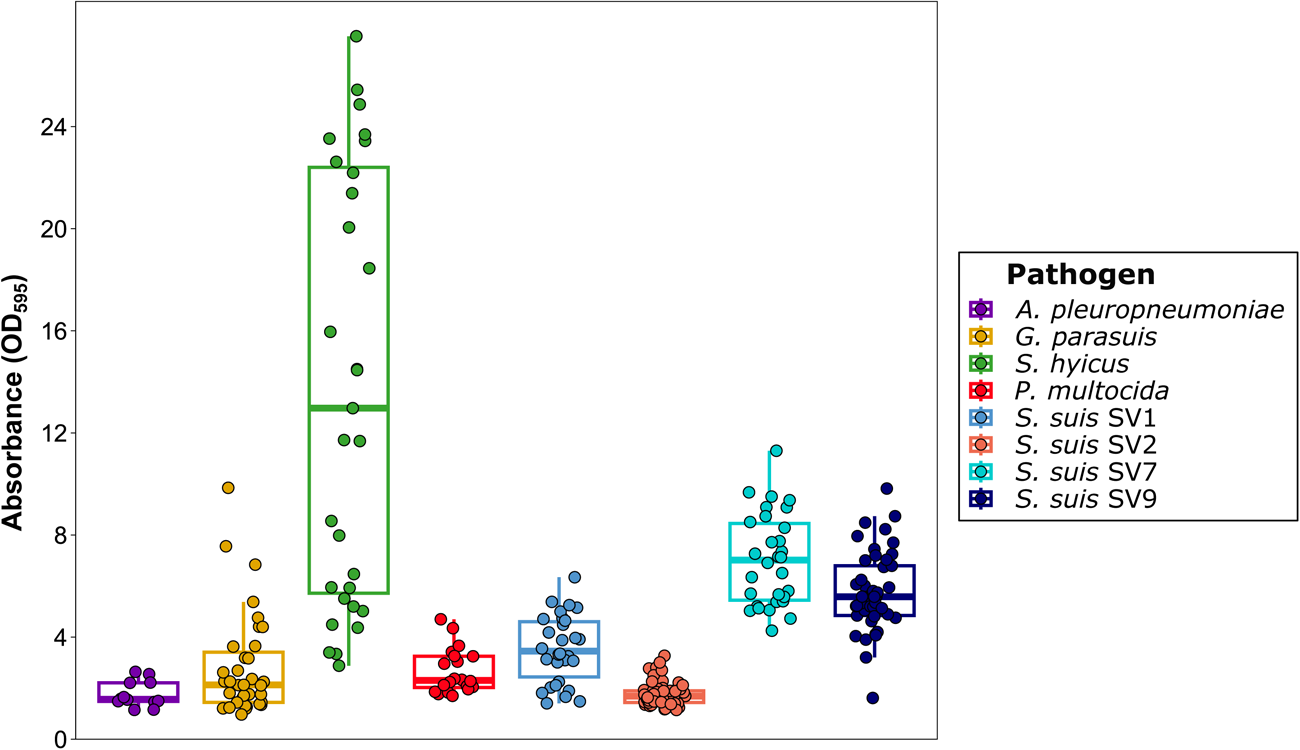
Biofilm formation of bacterial pathogens involved in the porcine respiratory disease complex (PRDC) and *Streptococcus suis* isolates belonging to the four main serovars (SVs). Boxplots illustrating biofilm formation for each isolate, quantified as the difference from the negative control (DC) in absorbance (OD_595_). Each isolate is represented by a dot with horizontal jitter for visibility. The horizontal box lines represent the first quartile, the median, and the third quartile. Whiskers extend to 1.5 times the interquartile range.

Interestingly, although *G. parasuis*, *P. multocida* and *A. pleuropneumoniae* produced significantly less biofilm than *S. suis* SV1, SV7 and SV9 (*p <* 0.05), the two formers had significantly higher biofilm production than *S. suis* SV2 (*p <* 0.05). A detailed summary of biofilm formation capacity comparisons among bacterial species and *S. suis* SVs is available in an additional file (see Additional file 5).

We further evaluated additional information of these bacterial pathogens, revealing that no significant associations were observed between biofilm production and VFs in *S. hyicus*, VFs and capsular type in *P. multocida*, SV in *A. pleuropneumoniae*, and virulence in *G. parasuis*. For coinfection studies, we assessed biofilm formation capacity of *S. suis* under microaerophilic conditions at 37°C for 48 h, replicating *A. pleuropneumoniae* and *G. parasuis* growth conditions, and compared it with standard growth conditions (37°C for 24 h under aerophilic conditions, optimal for *P. multocida* and *S. suis*). Given the clear differences between growth conditions, significant differences in biofilm formation capacity were observed, with a higher production under microaerophilic conditions (*p* < 0.05) also due to the supplementation with glucose (23). This finding does not interfere with further analyses, as coinfections were conducted individually for each pair of pathogens under the optimal growth conditions for the most fastidious microorganism.

### *In vitro* biofilm formation in coinfections between *Streptococcus suis* and clinically relevant swine bacterial pathogens

Notable differences were observed among *S. hyicus* isolates and *S. suis* SVs in coinfection (Figure 4A). Interestingly, those *S. hyicus* with the highest biofilm production, specifically H074, H086, H094, and H103, exhibited lower biofilm formation when coinfected with *S. suis*, regardless of the SV. In contrast, isolates with lower biofilm formation, such as H007, H026, H065, and H071, demonstrated a synergistic effect with *S. suis* coinfection, which was particularly remarkable when coinfected with SV9 and SV7. SEM revealed that the increased biofilm formation observed for *S. suis* in these coinfections was mainly determined by *S. hyicus*, with a reduced presence of *S. suis*, regardless of the *S. suis* SV (Figure 4B). When itemizing by *S. suis* SV, no significant differences were observed in the biofilm production of *S. hyicus* when coinfected with *S. suis*, but we observed a significant potentiation of biofilm formation in all *S. suis* SVs (*p* < 0.0001), nearly doubling the DC value in coinfections with SV1 and SV7, 3.2 times for SV9, and 4.2 times for SV2.

**Figure 4.**
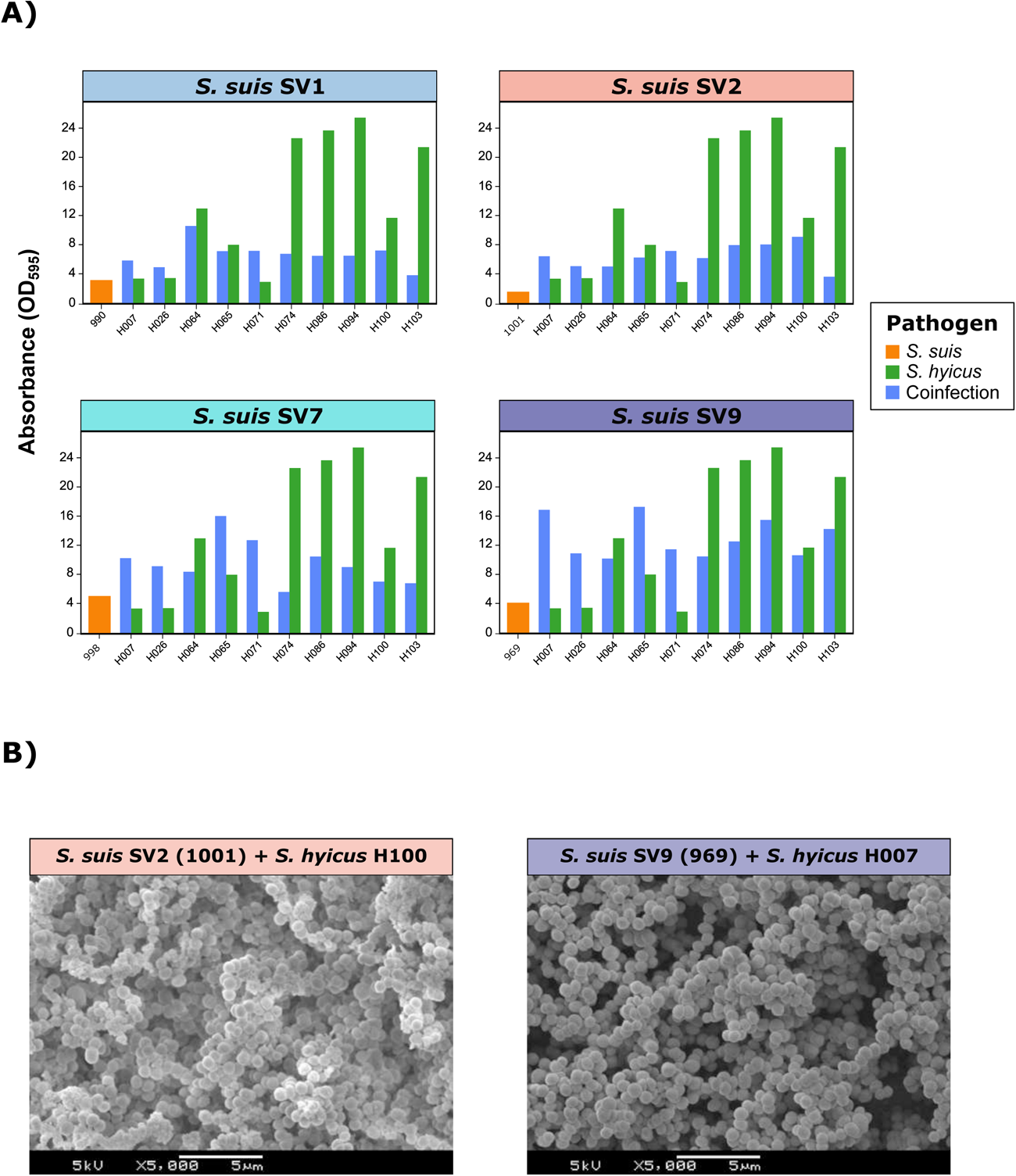
Biofilm formation in coinfections of *Streptococcus suis* and *Staphylococcus hyicus*. A) Bar plots showing the quantitative biofilm formation (expressed as absorbance OD_595_) of one representative *S. suis* isolate from each of the four main serovars (SVs) (orange), ten random *S. hyicus* isolates (green), and their coinfections (blue). B) Scanning electron microscope (SEM) images of biofilm formation in coinfections between *S. suis* SV2 (1001) and *S. hyicus* H100 (left) and *S. suis* SV9 (969) and *S. hyicus* H007 (right).

A synergistic biofilm production was demonstrated in coinfections between *S. suis* and certain *P. multocida* isolates, especially for PM179 with all SVs and, to a lesser extent, for PM182, except for SV2 (Figure 5A). Regarding the effect of the *S. suis* SV, an increase in biofilm formation in *P. multocida* isolates was noted when coinfected with SV7 (*p* < 0.01), nearly doubling the DC value, with a slightly higher contribution of *P. multocida* to the biofilm formation (Figure 5B). A synergistic effect of *P. multocida* coinfection was demonstrated for SV2, raising *S. suis* SV2 DC from 1.55 to 2.53 (*p <* 0.01). Notably, a potentiation was observed for both *P. multocida* (*p* < 0.01) and *S. suis* (*p* < 0.0001) in SV1 coinfection, increasing to a DC of 5.42 from 2.97 and 3.13, respectively (Figure 5B).

**Figure 5.**
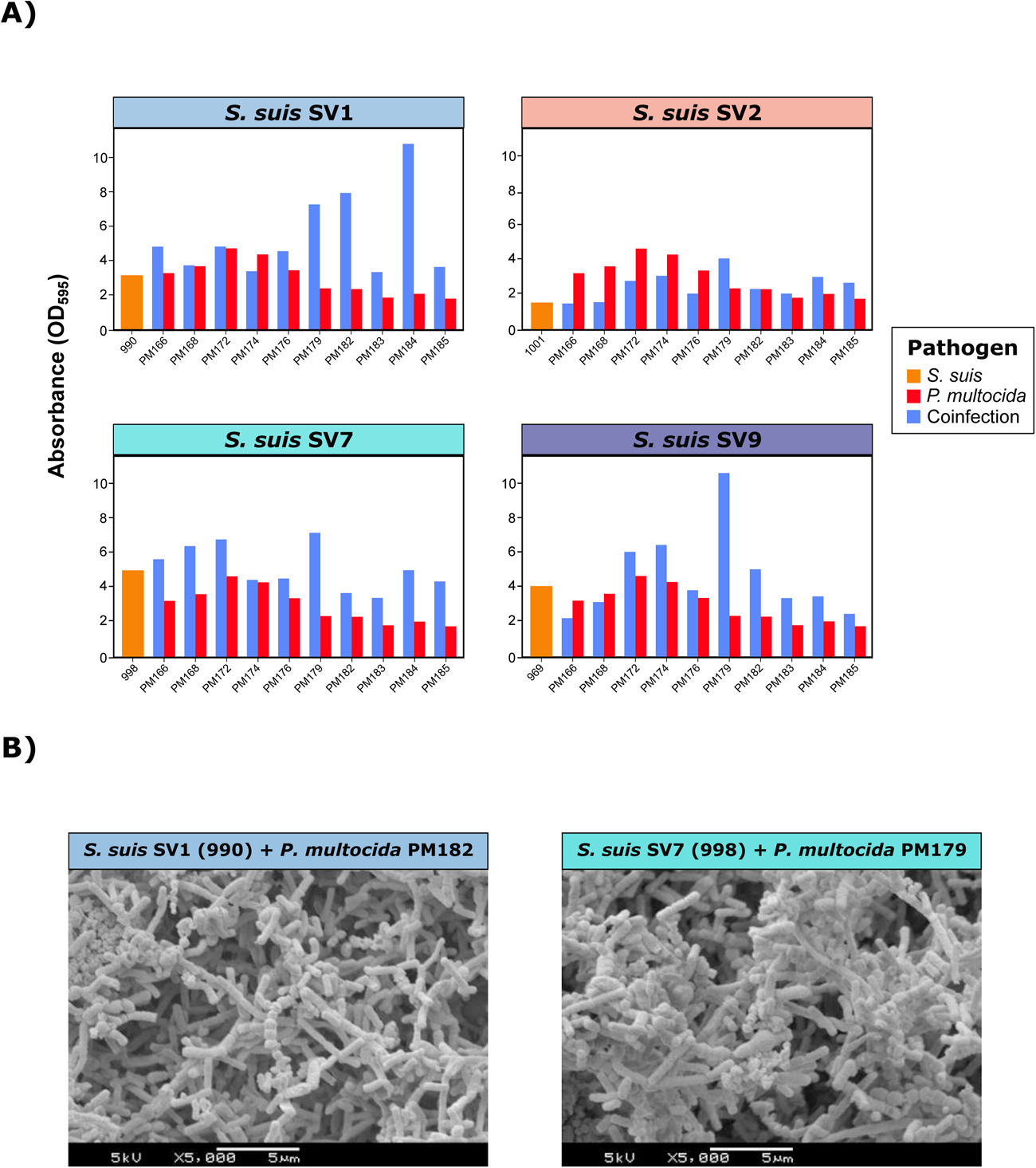
Biofilm formation in coinfections of *Streptococcus suis* and *Pasteurella multocida*. A) Bar plots showing the quantitative biofilm formation (expressed as absorbance OD_595_) of one representative *S. suis* isolate from each of the four main serovars (SVs) (orange), ten random *P. multocida* isolates (green), and their coinfections (blue). B) Scanning electron microscope (SEM) images of biofilm formation in coinfections between *S. suis* SV1 (990) and *P. multocida* PM182 (left) and *S. suis* SV7 (998) and *P. multocida* PM179 (right).

Coinfections between *G. parasuis* and *S. suis* revealed an overall significant reduction in biofilm formation for *S. suis* for SV2 (*p* < 0.05), SV7 (*p <* 0.001), and SV9 (*p* < 0.001) (Figure 6A). In contrast, *G. parasuis* increased its biofilm production in all SV coinfections, except for SV2, with a notable increase in SV1 (*p <* 0.001) and SV9 (*p <* 0.001), nearly 3.5 times the single *G. parasuis* DC average. Similar but more pronounced results were observed for *A. pleuropneumoniae* (Figure 7A). Coinfection increased biofilm formation in *A. pleuropneumoniae*, regardless of the *S. suis* SV (*p <* 0.05), particularly for SV9 (DC increase of 4.5 times), and SV7 and SV1 (DC increase of 2.6 times). Conversely, all *S. suis* SVs exhibited a significant reduction in biofilm production (*p <* 0.001). Remarkably, the only coinfection that increased biofilm formation was *S. suis* SV7 and *A. pleuropneumoniae* APP8. For both *G. parasuis* and *A. pleuropneumoniae* coinfections, SEM revealed that biofilm formation was mainly determined by *S. suis* (Figures 6B and 7B), with a reduced contribution of these pathogens to the biofilm matrix. All these findings demonstrate that the association between *S. suis* and other bacterial pathogens is not homogeneous, and substantial differences among SVs need to be considered.

**Figure 6.**
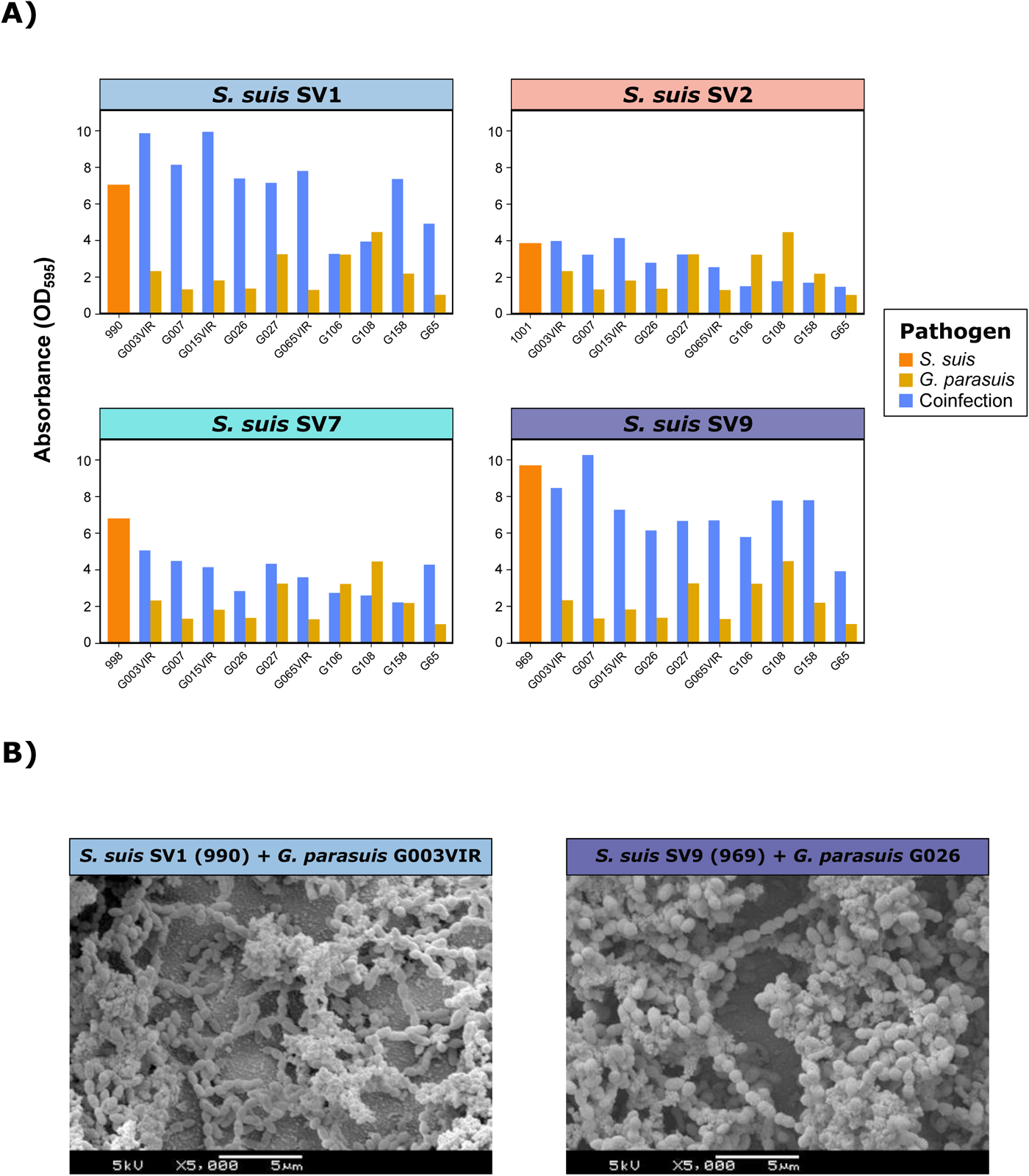
Biofilm formation in coinfections of *Streptococcus suis* and *Glaesserella parasuis*. A) Bar plots showing the quantitative biofilm formation (expressed as absorbance OD_595_) of one representative *S. suis* isolate from each of the four main serovars (SVs) (orange), ten random *G. parasuis* isolates (green), and their coinfections (blue). B) Scanning electron microscope (SEM) images of biofilm formation in coinfections between *S. suis* SV1 and *G. parasuis* G003 (left) and *S. suis* SV9 (969) and *G. parasuis* G026 (right).

**Figure 7.**
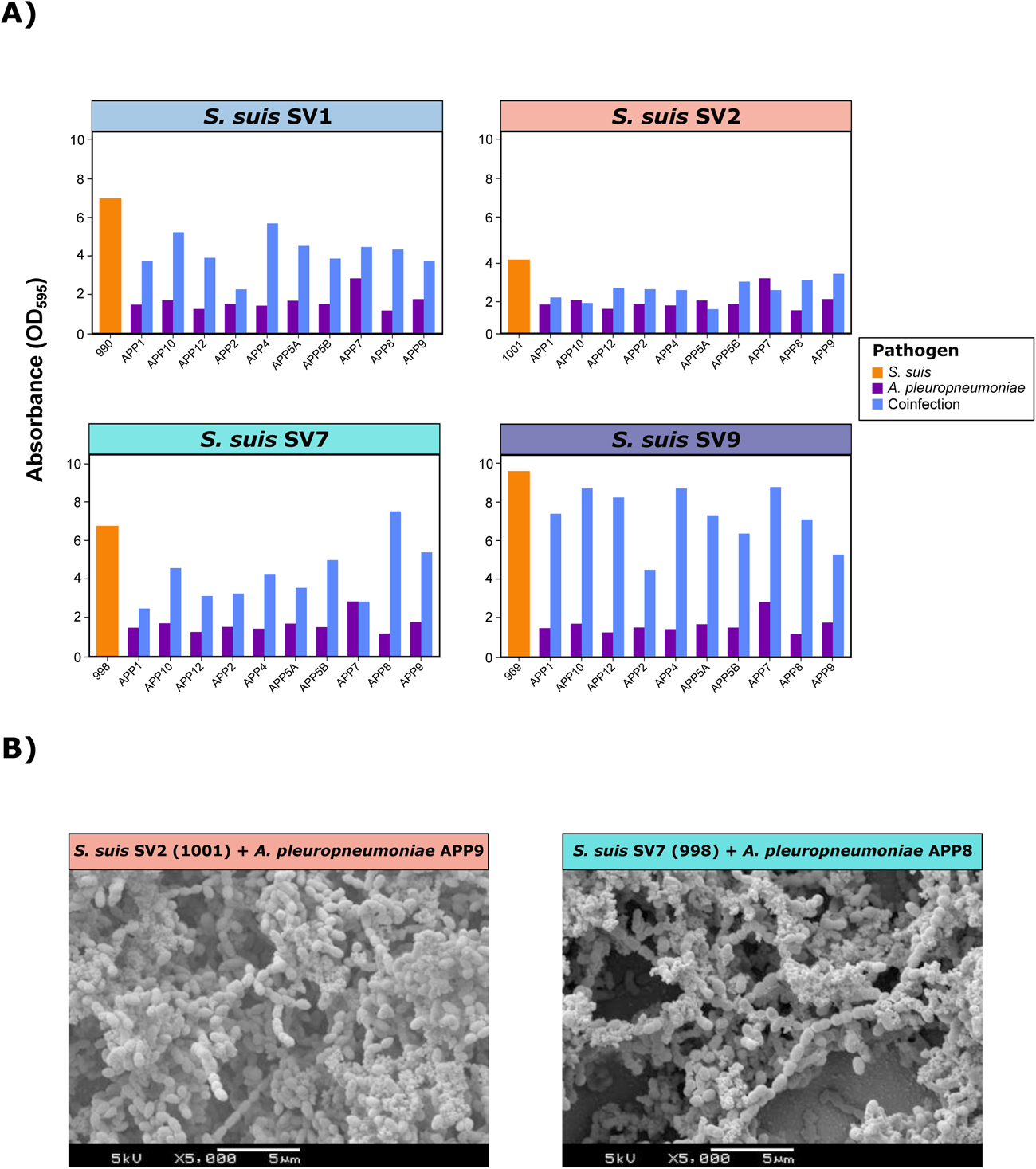
Biofilm formation in coinfections of *Streptococcus suis* and *Actinobacillus pleuropneumoniae*. A) Bar plots showing the quantitative biofilm formation (expressed as absorbance OD_595_) of one representative *S. suis* isolate from each of the four main serovars (SVs) (orange), ten random *A. pleuropneumoniae* isolates (green), and their coinfections (blue). B) Scanning electron microscope (SEM) images of biofilm formation in coinfections between *S. suis* SV2 (1001) and *A. pleuropneumoniae* APP9 (left) and *S. suis* SV7 (998) and *A. pleuropneumoniae* APP8 (right).

A detailed description of the significant interactions between *S. suis* SVs and bacterial pathogens, including the effects on both *S. suis* and the bacterial pathogens, along with DC averages and potential increases, is available in additional file (see Additional files 6 and 7).

## Discussion

*Streptococcus suis* is an opportunistic and zoonotic pathogen that naturally colonizes the respiratory tract in pigs (24). Several factors, such as bacterial or viral coinfections and environmental stressors, can cause *S. suis* to transition from a commensal to a pathogenic state (3). Its pathogenesis involves several niche environments, with multiple virulence mechanisms, including VFs and biofilm formation (6). Here, through extensive screening of virulence mechanisms in *S. suis* clinical isolates recovered from Spanish swine farms, we demonstrate that biofilm-forming ability is a significant pathogenic factor in certain *S. suis* SVs, particularly in those isolates harboring fewer virulence genes. Furthermore, *S. suis* interactions in biofilm formation with bacteria involved in the PRDC clearly vary among SVs and pathogens.

The multifactorial nature of *S. suis* pathogenicity was evident in our study, which revealed a substantial diversity in the frequency of VFs among isolates, with 27 distinct gene patterns observed. Approximately one-third of the isolates (33.8%) carried all five evaluated VFs. Among these, *mrp*, *epf*, and *sly* genes, which are frequently associated with virulence (25), were simultaneously present in 37.9% of the clinical *S. suis* isolates. Additionally, 86.7% and 66.7% of them harbored one or two of these genes, respectively. This finding aligns with previous studies suggesting that their absence is more commonly associated with *S. suis* isolates from healthy or carrier pigs in Europe and Asia (26–28). Nonetheless, the absence of one or more of these VFs does not necessarily correlate with a lack of virulence (5, 29). For instance, *sly* is typically absent in isolates from North America, but these isolates do not exhibit reduced virulence compared to *sly*-producing *S. suis* (30). The complexity of VF associations is further highlighted by the relationship between VFs and *S. suis* SVs, as recently described (31). It is particularly remarkable for SV7, in which a negative association with the presence of *epf* and *sly* was observed, consistent with a previous study on *S. suis* SV7 clinical isolates from pig farms in Germany (32). These findings underscore the importance of considering other mechanisms, such as biofilm formation, when evaluating *S. suis* pathogenicity.

Biofilm formation is an essential pathogenic mechanism in *S. suis*, enabling its establishment in pig tissues, as most isolates can form biofilms (7). Indeed, the development of bacterial meningitis is strongly associated with biofilm formation (33). However, differences exist among strains. In this study, we demonstrate that *in vitro* biofilm formation is deeply influenced by the *S. suis* SV. A previous investigation revealed differences in biofilm-forming ability between SV2 and SV9 isolates (23), and our wide-range assessment of 16 different Spanish *S. suis* SVs expands on this finding.

*S. suis* SV2 was the only SV categorized as low biofilm-forming. Despite its prevalence in swine infections (34) and its role as a primary lineage in human infections worldwide (35), our study shows that, interestingly, biofilm formation is not an essential pathogenic factor for SV2, contrasting with other SVs. For instance, *S. suis* SV9, an important and prevalent SV causing invasive disease in pigs in Europe (36), demonstrated strong biofilm formation regardless of its virulence gene arsenal.

Reduced virulence has been described as an important characteristic of biofilm infection in *S. suis* (37). Our research indicates an overall significant reduction in biofilm formation among *S. suis* isolates harboring *epf*, *mrp*, and *sly* genes. Previous studies have shown a differential expression of virulence genes under planktonic and biofilm conditions (37), which could explain the presence of virulent strains in the host respiratory tract as commensals. In addition, *S. suis* in a biofilm-state is less likely to trigger the immune system (38). Although *luxS* is involved in the LuxS/AI-2 quorum sensing (QS) system, a crucial regulatory network influencing biofilm formation (39), no significant differences in biofilm formation were observed when evaluating isolates harboring *luxS*. This could be due to its high prevalence in clinical *S. suis*, underscoring its importance as a virulence determinant (40). Interestingly, when analyzing the association between biofilm formation and VFs by SV, we found that the overall differences were reduced. Notably, *S. suis* SV1 isolates carrying the *epf* gene exhibited significantly higher biofilm formation, aligning with studies suggesting that highly pathogenic strains may exhibit strong biofilm formation (37). These differences likely depend on specific gene expression patterns in biofilm state rather than the mere presence of the genes. Therefore, further investigations are necessary to evaluate gene expression changes between planktonic and biofilm-forming cells among SVs with varying biofilm-forming abilities.

Other bacterial pathogens have also been shown to form biofilms within the PRDC (41), and their potential role in coinfections with *S. suis* should be considered. Here, we demonstrated the *in vitro* biofilm formation abilities of primary (*A. pleuropneumoniae*) and secondary (*G. parasuis* and *P. multocida*) PRDC pathogens recovered from Spanish swine farms. However, the biofilms formed by these bacterial pathogens were generally less robust than those formed by most *S. suis* SVs, except for *S. suis* SV2. The low to medium biofilm formation observed in *A. pleuropneumoniae* aligns with previous studies showing the biofilm-forming ability of most field isolates (42), since this pathogen is known to form biofilms in lungs (43).

For *G. parasuis*, we observed a wide range of biofilm-forming abilities among our isolates, regardless of their virulence, with an overall medium biofilm formation.

Recently, 76 genes have been identified as potentially involved in *G. parasuis* biofilm formation. Nonetheless, differences were observed among isolates, even within the same SV, likely due to its open pangenome and variations in the accessory genome (44).

*P. multocida* was identified as an intermediate biofilm producer compared to other PRDC pathogens, with 65% of clinical isolates characterized as mid-level biofilm producers, consistent with recent findings (45). An inverse association between capsular polysaccharide production and biofilm formation has been described, with encapsulated *P. multocida* isolates, presumed to be more virulent, producing less biofilm than those with reduced capsular polysaccharide (46). However, we could not corroborate this finding due to the limited *P. multocida* isolates used in the study and the fact that all of them were clinical and produced capsular polysaccharide. Since *P. multocida* was not the primary focus of this study, our results did not explore these differences in depth, and, hence, further investigations are needed.

Despite *S. hyicus*, the causative agent of exudative epidermitis, is not regarded as a constituent of the PRDC, recent studies have described its potential role in swine respiratory disease cases (47, 48), leading us to its consideration in the biofilm persistence of bacterial pathogens involved in the PRDC. In this study, the strong biofilm formation capacity of *S. hyicus* could be proven, with more than 90% of the isolates showing robust biofilm formation, several of which were well above the threshold to be considered strong biofilm formers. To the best of our knowledge, this is the first report specifically addressing the biofilm formation of *S. hyicus*, although previous studies have evaluated its formation within sets of coagulase-negative staphylococci, yielding disparate results (49). The strong biofilm formation observed in this study could serve as a starting point for future studies aimed at understanding the mechanisms of *S. hyicus* biofilm formation and its impact on the microbial environment in the respiratory tract.

Given the high prevalence of bacterial coinfections in the PRDC (50), mixed biofilms may be common and contribute to enhanced bacterial survival through interspecific competition, communication, and cooperation (8). However, potential interactions involving *S. suis* have been scarcely studied, with a primary focus on *S. suis* SV2, which was shown to be the lowest biofilm former among *S. suis* SVs. This research reveals that the contribution of each pathogen in *in vitro* biofilm formation differs depending on the bacterial pathogen and the *S. suis* SV involved. Notably, this is the first report to show that biofilm formation was stronger in mixed infections with *S. suis* and *P. multocida* than in single infections for both microorganisms, with a clear presence of both pathogens in the biofilm matrix. Additionally, we demonstrate an overall promotion of biofilm formation for *G. parasuis* and *A. pleuropneumoniae* when coinfected with *S. suis*, although it was mainly determined by the presence of *S. suis*. Thus, *S. suis* could contribute to the persistence of these pathogens integrated in the biofilm matrix. These findings are consistent with previous studies that analyzed the role of *S. suis* in the persistence of both *G. parasuis* (51) and *A. pleuropneumoniae* (52) in mixed biofilms with *S. suis* SV2. In the case of *A. pleropneumoniae*, biofilm growth was promoted under hostile conditions, such as the absence of NAD, when coinfected with *S. suis* (53), revealing that mixed infections may be more difficult to eradicate. In contrast, *S. suis* coinfection with *S. hyicus* revealed that the latter was the main determinant in the biofilm matrix, highlighting its potential relevance in the persistence of *S. suis* in the respiratory tract, since both are frequently present in tonsils (54). Advancing the knowledge in the role of biofilm formation in respiratory mixed infections will contribute to the establishment of optimal control measures for the PRDC, a syndrome that causes significant economic losses in pig production worldwide (55).

## Conclusions

This study highlights the heterogeneity in virulence factors and *in vitro* biofilm formation among *S. suis* clinical isolates from Spanish swine farms, particularly influenced by SV variations. Our findings state that while some *S. suis* SVs, such as SV2, show low biofilm-forming abilities, others like SV1, SV7 or SV9, exhibit robust biofilm formation, independent of their virulence gene arsenal. Additionally, the present study underscores the complexity of mixed biofilm formation in coinfections, revealing heterogeneous biofilm production in interactions between *S. suis* and other primary or secondary PRDC bacterial pathogens, such as *P. multocida*, *G. parasuis*, or *A. pleuropneumoniae*. Remarkably, *S. hyicus*, typically not associated with PRDC, displayed strong biofilm formation, suggesting its potential role in *S. suis* persistence in the upper respiratory tract. These insights pave the way for more detailed investigations into the mechanisms underlying biofilm formation and maintenance in PRDC-associated pathogens, ultimately contributing to the development of effective control measures to mitigate its economic impact in the swine industry.

## List of abbreviations

EF: extracellular factor Exh: exfoliative toxin
FBS: fetal bovine serum
GAPDH: glyceraldehyde-3-phosphate dehydrogenase
LuxS: S-ribosylhomocysteinase
MRP: muramidase-released protein
NAD: Nicotinamide adenine dinucleotide
OD: optical density
PBS: phosphate-buffered saline
PCA: principal component analyisis
PCR: polymerase chain reaction
PRDC: Porcine respiratory disease complex
SEM: scanning electron microscopy
SV: Serovar
Tbp: Transferring-binding protein
THB: Todd-Hewitt broth
VF: virulence factor

## Declarations

### Ethics approval and consent to participate

Not applicable.

### Consent for publication

Not applicable.

### Availability of data and material

Not applicable.

### Authors’ contributions

Study design was performed by SMM, CBGM and OMA. Laboratory analyses were performed by RMP with support of AGF, MP, and MDG. Statistical analyses were performed by OMA. SMM, CBGM, and OMA provided technical and scientific support on the analysis. RMP, OMA, CBGM and SMM participated in the manuscript writing or contributed to its revision. All authors revised the manuscript and approved the final version.

## Acknowledgments

We acknowledge the excellent technical assistance provided by María Mediavilla and Vanessa Acebes. We would also like to thank the contribution of University of León’s microscopy services and personnel, for their predisposition and assistance.

